## Supplemental Materials for "Knockout of *myoc* reveals the role of myocilin in zebrafish sex determination associated with Wnt signalling downregulation"

**Table S1.** Primers sequences used in RT-qPCR.

| Gene | Forward primer (5'-3') | Reverse primer (5'-3') | Amplicon size (pb) | Reference |
|---|---|---|---|---|
| *amh* | GACCTTGAGGAGCCTCGTTT | GTACTTTTGCTCTGAGGCAGG | 128 | |
| *ctnnbip1* | CTGTCGGGATGTGACCCCGG | CTCCTGACGCACCGCTCTCC | 106 | [1] |
| *cyp11a1* | TCCTGTAGCAATGAGTCTTCC | GATGATCACGACCCATAGCA | 116 | |
| *dkk1a* | CCTGTCAAAACCACAGCAGA | GTCAGCACCGGCTTACAGAT | 126 | |
| *dmrt1* | TGCCCAGGTGGCGTTACGG | CGGGTGATGGCGGTCCTGAG | 149 | [1] |
| *dvl3a* | TGCCCATCCCTGCCGAAAGG | TGACCACCCCAAAGTCATCGTCC | 108 | [1] |
| *ef1α* | CTGGAGGCCAGCTCAAACAT | ATCAAGAAGAGTAGTACCGCTAGCATTAC | 87 | |
| *grik3* | AGTATGTGACGCGCAGGAA | CCCAGGATAGCAATGGTGA | 124 | |
| *irg1l* | CAGAGCGTACAGCCAAAGAA | CCCAACTCCAATGCTGTCTAA | 81 | |
| *lef1* | CGAGGGAGACCCGCACAAGG | GGGACTGTCTAGCTGCGTCGTG | 146 | [1] |
| *myoc* | CAAAAAGACAACAGTTCAGACC | GACCACAGCCGTCAACTC | 149 | |
| *plpp4* | CCCGCATCTGTGACTACAAA | GTGGCAGTCTATGTGCAGGA | 122 | |
| *stAR* | AGTGGAACCCCAATGTCAAG | TCTTGGGCCTACCACATTTC | 110 | |
| *sycp3* | AGCGGATCTGACGAAGACACGAG | ATGTCCGCACCAAATCTTTCCAGC | 149 | [1] |
| *tuba7l* | CACACTGCTCTCTGGACTTTG | GGTGCCCAAGGATGTCAAC | 159 | [2] |

**Table S2.** Top-50 downregulated genes identified by the transcriptomic study in the four comparisons (KO1 vs. WT1, KO1 vs. WT2, KO2 vs. WT1 and KO2 vs. WT2). Differential expression threshold: foldchange <-2; read count>20. p-value<0.05.

| Gene symbol | Gene ID | Identificaction | Fold change (KO vs WT) | P-value |
|---|---|---|---|---|
| *plpp4* | 100334794 | phospholipid phosphatase 4 | -219.7 | 5.1E-04 |
| *grik3* | 100334689 | glutamate receptor ionotropic, kainate 3 | -189.5 | 1.8E-04 |
| *cth1* | 30114 | cysteine three histidine 1 | -61.4 | 6.7E-49 |
| *birc5b* | 246726 | baculoviral IAP repeat containing 5b | -59.6 | 1.9E-36 |
| *Cldnd* | 335628 | claudin domain containing 1a | -57.9 | 4.1E-08 |
| *bmp15* | 334183 | bone morphogenetic protein 15 | -55.4 | 5.5E-16 |
| *hsd3b2* | 373131 | hydroxy-delta-5-steroid dehydrogenase, 3 beta- and steroid delta-isomerase 2 | -53.3 | 4.4E-41 |
| *si:dkey-152b24.6* | 100002310 | si:dkey-152b24.6 | -52.1 | 3.1E-16 |
| *LOC100332229* | 100332229 | histone H2A, sperm-like | -51.1 | 7.8E-17 |
| *btg4* | 378946 | B-cell translocation gene 4 | -49.7 | 2.2E-40 |
| *zgc:55413* | 406830 | zgc:55413 | -49.2 | 1.0E-45 |
| *si:ch211-286b5.4* | 556026 | si:ch211-286b5.4 | -47.3 | 3.9E-44 |
| *si:ch211-103b1.2* | 556271 | si:ch211-103b1.2 | -46.6 | 1.9E-49 |
| *tuba4l* | 327377 | tubulin, alpha 4 like | -46.6 | 8.7E-48 |
| *Retsatl* | 677660 | retinol saturase (all-trans-retinol 13,14-reductase) like | -45.9 | 1.2E-46 |
| *hyal6* | 791189 | hyaluronoglucosaminidase 6 | -45.5 | 5.4E-22 |
| *zgc:77118* | 405845 | zgc:77118 | -45.2 | 6.2E-43 |
| *zgc:171517* | 563255 | zgc:171517 | -44.7 | 9.9E-10 |
| *Cldng* | 81586 | claudin g | -43.9 | 7.2E-46 |
| *ca15b* | 791844 | carbonic anhydrase XVb | -42.3 | 1.1E-43 |
| *LOC103911725* | 103911725 | molybdopterin synthase catalytic subunit-like | -42.2 | 2.6E-15 |
| *cpeb1b* | 30702 | cytoplasmic polyadenylation element binding protein 1b | -40.2 | 4.5E-14 |
| *pabpc1l* | 327625 | poly(A) binding protein, cytoplasmic 1-like | -38.9 | 9.9E-48 |
| *mcm6l* | 564982 | MCM6 minichromosome maintenance deficient 6, like | -38.7 | 1.8E-41 |
| *gtf2f2b* | 100148942 | general transcription factor IIF, polypeptide 2b | -38.7 | 7.0E-31 |
| *zgc:66024* | 393657 | zgc:66024 | -37.3 | 3.8E-45 |
| *gdf3* | 30125 | growth differentiation factor 3 | -35.1 | 3.7E-48 |
| *ccna1* | 404206 | cyclin A1 | -34.2 | 4.4E-45 |
| *map1lc3c* | 393268 | microtubule-associated protein 1 light chain 3 gamma | -33.6 | 4.8E-44 |
| *st6galnac1.2* | 799868 | ST6 (alpha-N-acetyl-neuraminyl-2,3-beta-galactosyl-1,3)-N-acetylgalactosaminide alpha-2,6-sialyltransferase 1, tandem duplicate 2 | -32.9 | 1.4E-39 |
| *Nots* | 114367 | Nothepsin | -32.9 | 8.0E-19 |
| *ccnb2* | 368316 | cyclin B2 | -31.7 | 1.3E-45 |
| *si:dkey-37o8.1* | 407641 | si:dkey-37o8,1 | -30.2 | 2.5E-42 |
| *si:busm1-194e12.12* | 368615 | si:busm1-194e12.12 | -30.2 | 1.9E-53 |
| *gdf9* | 497643 | growth differentiation factor 9 | -30.1 | 1.2E-43 |

**Table S2.** Top-50 downregulated genes identified by the transcriptomic study in the four comparisons (KO1 vs. WT1, KO1 vs. WT2, KO2 vs. WT1 and KO2 vs. WT2). Differential expression threshold: foldchange <-2; read count>20. p-value<0.05.

| Gene symbol | Gene ID | Identificaction | Fold change (KO vs WT) | P-value |
|---|---|---|---|---|
| *si:busm1-48c11.3* | 368614 | si:busm1-48c11.3 | -28.3 | 4.2E-41 |
| *Mos* | 795517 | v-mos Moloney murine sarcoma viral oncogene homolog | -27.9 | 3.2E-38 |
| *pabpn1l* | 796580 | poly(A) binding protein, nuclear 1-like (cytoplasmic) | -27.5 | 4.4E-38 |
| *polr2ea* | 100330570 | polymerase (RNA) II (DNA directed) polypeptide E, a | -26.9 | 3.3E-32 |
| *siva1* | 571845 | SIVA1, apoptosis-inducing factor | -26.7 | 8.1E-42 |
| *mcm3l* | 100147915 | MCM3 minichromosome maintenance deficient 3 (S, cerevisiae), like | -26.2 | 3.3E-46 |
| *polr3g* | 449786 | polymerase (RNA) III (DNA directed) polypeptide G like b | -22.7 | 1.7E-03 |
| *cyp11a1* | 80374 | cytochrome P450, family 11, subfamily A, polypeptide 1 | -22.3 | 5.6E-40 |
| *ccnb1* | 562825 | cyclin B1 interacting protein 1 | -20.7 | 2.4E-04 |
| *fbxo43* | 393403 | F-box protein 43 | -18.8 | 1.3E-40 |
| *wee2* | 327471 | WEE1 homolog 2 (S, pombe) | -17.9 | 4.2E-41 |
| *rdh10b* | 378722 | retinol dehydrogenase 10b | -17.0 | 1.6E-13 |
| *tph1a* | 352943 | tryptophan hydroxylase 1 (tryptophan 5-monooxygenase) a | -15.7 | 4.4E-20 |
| *st6gal1* | 445376 | ST6 beta-galactosamide alpha-2,6-sialyltranferase 1 | -13.0 | 1.0E-10 |
| *rag1* | 30663 | recombination activating gene 1 | -8.1 | 1.1E-26 |

**Table S3.** Top-50 upregulated genes identified by the transcriptomic study in the four comparisons (KO1 vs. WT1, KO1 vs. WT2, KO2 vs. WT1 and KO2 vs. WT2). Differential expression threshold: foldchange >2; read count >20. p-value<0.05.

| Gene symbol | Gene ID | Identification | Fold change (KO vs WT) | P-value |
|---|---|---|---|---|
| *tuba7l* | 431777 | tubulin, alpha 7 like | 49.9 | 7.2E-25 |
| *irg1l* | 562007 | immunoresponsive gene 1, like | 35.4 | 1.5E-23 |
| *smc1b* | 797354 | structural maintenance of chromosomes 1B | 20.0 | 2.4E-11 |
| *sat2a* | 494055 | spermidine/spermine N1-acetyltransferase family member 2a | 18.8 | 6.7E-31 |
| *Ddc* | 406651 | dopa decarboxylase | 15.9 | 1.4E-39 |
| *sult2st3* | 777792 | sulfotransferase family 2, cytosolic sulfotransferase 3 | 15.2 | 2.4E-16 |
| *Hamp* | 402837 | hepcidin antimicrobial peptide | 15.2 | 8.5E-27 |
| *hsd17b7* | 768185 | hydroxysteroid (17-beta) dehydrogenase 7 | 14.2 | 2.1E-22 |
| *cremb* | 550357 | cAMP responsive element modulator b | 12.5 | 3.5E-13 |
| *sycp3* | 678602 | synaptonemal complex protein 3 | 12.1 | 2.6E-21 |
| *zgc:92137* | 445049 | zgc:92137 | 11.9 | 1.3E-27 |
| *cyp8b2* | 100004274 | cytochrome P450, family 8, subfamily B, polypeptide 2 | 11.7 | 1.6E-04 |
| *eno4* | 558487 | enolase family member 4 | 11.5 | 2.0E-17 |
| *Cthl* | 445818 | cystathionase (cystathionine gamma-lyase), like | 11.3 | 9.0E-46 |
| *gstk4* | 678541 | glutathione S-transferase kappa 4 | 11.2 | 7.7E-08 |
| *soat2* | 564868 | sterol O-acyltransferase 2 | 11.2 | 1.1E-09 |
| *si:busm1-194e12.8* | 368816 | si:busm1-194e12.8 | 10.9 | 1.2E-23 |
| *ak7b* | 504040 | adenylate kinase 7b | 10.0 | 4.5E-05 |
| *cyp11c1* | 791124 | cytochrome P450, family 11, subfamily C, polypeptide 1 | 10.0. | 6.1E-13 |
| *galnt8a.2* | 568500 | polypeptide N-acetylgalactosaminyltransferase 8a, tandem duplicate 2 | 9.2 | 4.4E-21 |
| *si:ch211-199m3.4* | 557708 | si:ch211-199m3,4, transcript variant X1 | 9.0 | 1.3E-17 |
| *rad9b* | 563903 | RAD9 checkpoint clamp component B | 8.6 | 6.5E-04 |
| *zgc:195023* | 567953 | zgc:195023 | 7.9 | 6.2E-15 |
| *nos2a* | 404036 | nitric oxide synthase 2a, inducible | 7.5 | 1.0E-28 |
| *spam1* | 791203 | sperm adhesion molecule 1 | 7.5 | 9.6E-20 |
| *trnG* | 140510 | tRNA-Gly | 7.5 | 1.9E-03 |
| *amh* | 493624 | anti-Mullerian hormone | 6.9 | 1.6E-20 |
| *tnnc1a* | 353247 | troponin C type 1a (slow) | 6.8 | 4.9E-21 |
| *ugt5a1* | 641479 | UDP glucuronosyltransferase 5 family, polypeptide A1 | 6.1 | 1.9E-04 |
| *LOC100007686* | 100007686 | phosphomannomutase 1-like | 6.1 | 6.6E-06 |
| *cyp46a1.2* | 393433 | cytochrome P450, family 46, subfamily A, polypeptide 1, tandem duplicate 2 | 6.1 | 6.6E-06 |
| *gch1* | 100192219 | GTP cyclohydrolase 1 | 6.0. | 6.1E-06 |
| *sqlea* | 799528 | squalene epoxidase a | 5.9 | 5.2E-11 |
| *upp2* | 393113 | uridine phosphorylase 2 | 5.9 | 8.1E-23 |
| *pcyt1ba* | 100001552 | phosphate cytidylyltransferase 1, choline, beta a | 5.9 | 3.8E-09 |
| *LOC101885031* | 101885031 | laminin subunit beta-3-like | 5.8 | 9.2E-19 |

**Table S3.** Top-50 upregulated genes identified by the transcriptomic study in the four comparisons (KO1 vs. WT1, KO1 vs. WT2, KO2 vs. WT1 and KO2 vs. WT2). Differential expression threshold: foldchange >2; read count >20. p-value<0.05.

| Gene symbol | Gene ID | Identification | Fold change (KO vs WT) | P-value |
|---|---|---|---|---|
| *pnp6* | 402953 | purine nucleoside phosphorylase 6 | 5.7 | 3.9E-07 |
| *trnL1* | 140530 | tRNA-Leu | 5.7 | 2.8E-17 |
| *haao* | 492518 | 3-hydroxyanthranilate 3,4-dioxygenase | 5.5 | 1.3E-05 |
| *itga2.2* | 100536533 | integrin, alpha 2 (CD49B, alpha 2 subunit of VLA-2 receptor), tandem duplicate 2, transcript variant X2 | 5.4 | 8.0E-12 |
| *miox* | 571850 | myo-inositol oxygenase | 5.1 | 5.8E-15 |
| *cldn10b* | 100004456 | claudin 10b | 5.1 | 1.1E-14 |
| *dkk1a* | 799377 | dickkopf WNT signaling pathway inhibitor 1a | 4.6 | 6.2E-11 |
| *ca9* | 566612 | carbonic anhydrase IX, transcript variant X4 | 4.5 | 3.4E-13 |
| *casp8l2* | 557302 | caspase 8, apoptosis-related cysteine peptidase, like 2 | 4.5 | 2.6E-07 |
| *si:dkey-33b17.1* | 795595 | si:dkey-33b17.1 | 4.5 | 5.5E-08 |
| *inhbab* | 553816 | inhibin, beta Ab | 4.3 | 2.4E-08 |
| *ndufa4l2a* | 100003066 | NADH dehydrogenase (ubiquinone) 1 alpha subcomplex, 4-like 2a | 4.2 | 9.5E-11 |
| *crb3a* | 724017 | crumbs homolog 3a | 3.9 | 6.3E-14 |
| *mhc2dab* | 30762 | major histocompatibility complex class II DAB gene | 3.8 | 1.4E-13 |

**Table S4.** ShinyGO ontological analysis of the top-50 upregulated genes in the transcriptomic study. p-value <0.05.

| N | High level GO category | Genes |
|---|---|---|
| 13 | Biosynthetic process | *nos2a, cthl, cyp11c1, gch1, sqlea, hsd17b7, galnt8a,2, ak7b, upp2, pcyt1ba, eno4, haao, cremb* |
| 7 | Developmental process | *sycp3, inhbab, dkk1a, amh, tnnc1a, ddc, crb3a* |
| 7 | Anatomical structure development | *sycp3, inhbab, dkk1a, amh, tnnc1a, ddc, crb3a* |
| 7 | Regulation of molecular function | *nos2a, miox, casp8l2, dkk1a, amh, inhbab, zgc:195023, hamp* |
| 6 | Catabolic process | *Nos2a, miox, cyp46a1,2, upp2, eno4, haao* |
| 5 | Response to stress | *Irgl1, rad9b, nos2a, hamp, dkk1a* |
| 5 | Regulation of signaling | *dkk1a, inhbab, amh, zgc:195023, hamp* |
| 5 | Response to chemical | *nos2a, cyp11c1, sult2st3, gstk4, hamp* |
| 5 | Regulation of response to stimulus | *dkk1a, inhbab, amh, zgc:195023, hamp* |
| 5 | Multi-organism process | *sycp3, nos2a, hamp, amh, irg1l* |
| 5 | Regulation of biological quality | *nos2a, cyp11c1, amh, zgc:195023, hamp* |
| 4 | System process | *tnnc1a, nos2a, amh, zgc:195023* |
| 4 | Response to endogenous stimulus | *nos2a, cyp11c1, sult2st3, hamp* |
| 4 | Regulation of metabolic process | *inhbab, nos2a, casp8l2, cremb* |
| 3 | Response to external stimulus | *nos2a, hamp, irg1l* |
| 3 | Response to biotic stimulus | *nos2a, hamp, irg1l* |
| 3 | Cellular component organization | *tuba7l, smc1b, crb3a* |
| 3 | Positive regulation of biological process | *inhbab, nos2a, casp8l2* |
| 3 | Negative regulation of biological process | *dkk1a, rad9b, amh* |
| 3 | Localization | *soat2, amh, ndufa4l2a* |
| 3 | Establishment of localization | *soat2, amh, ndufa4l2a* |
| 3 | Response to other organism | *nos2a, hamp, irg1l* |
| 3 | Cellular component organization or biogenesis | *tuba7l, smc1b, crb3a* |
| 2 | Reproduction | *sycp3, amh* |
| 2 | Immune system process | *mhc2dab, si:busm1-194E12,8* |
| 2 | Developmental process involved in reproduction | *sycp3, amh* |
| 2 | Immune response | *mhc2dab, si:busm1-194E12,8* |
| 2 | Antigen processing and presentation | *mhc2dab, si:busm1-194E12,8* |
| 2 | Sexual reproduction | *sycp3, amh* |
| 2 | Reproductive process | *sycp3, amh* |
| 2 | Multicellular organism reproduction | *sycp3, amh* |
| 2 | Macromolecule localization | *sycp3, amh* |
| 2 | Multi-organism reproductive process | *sycp3, amh* |
| 2 | Multicellular organismal reproductive process | *sycp3, amh* |
| 2 | Regulation of multicellular organismal process | *sycp3, amh* |

**Table S5.** Ontological analysis of the top-50 downregulated genes in the transcriptomic study. p-value <0.05.

| N | High level GO category | Genes |
|---|---|---|
| 17 | Biosynthetic process | *cyp11a1, gdf9, gtf2f2b, cpeb1b, hsd3b2, bmp15, gdf3, cth1, si:dkey-37o8,1, zgc:171517, rdh10b, polr3g, tph1a, mcm6l, mcm3l, st6galnac1,2, st6gal1* |
| 12 | Positive regulation of biological process | *gdf9, gtf2f2b, ccnb2, bmp15, gdf3, ccna1, ccnb1, cth1, mos, cyp11a1, birc15b, cpeb1b* |
| 11 | Cellular component organization | *wee2, ccnb2, ccna1, ccnb1, birc5b, mos, tuba4l, map1lc3c, fbxo43, mcm6l, cyp11a1* |
| 11 | Cellular component organization or biogenesis | *wee2, ccnb2, ccna1, ccnb1, birc5b, mos, tuba4l, map1lc3c, fbxo43, mcm6l, cyp11a1* |
| 10 | Regulation of metabolic process | *gdf9, gtf2f2b, cpeb1b, ccnb2, bmp15, gdf3, ccna1, ccnb1, cth1, mos* |
| 9 | Developmental process | *gdf9, bmp15, gdf3, ccnb1, rag1, tph1a, cyp11a1, hsd3b2, birc5b* |
| 9 | Anatomical structure development | *gdf9, bmp15, gdf3, ccnb1, rag1, tph1a, cyp11a1, hsd3b2, birc5b* |
| 8 | Reproduction | *wee2, mos, fbxo43, bmp15, ccnb1, ca15b, gdf9, birc5b* |
| 8 | Reproductive process | *wee2, mos, fbxo43, bmp15, ccnb1, ca15b, gdf9, birc5b* |
| 8 | Negative regulation of biological process | *cpeb1b, wee2, btg4, cth1, mos, fbxo43, bmp15, birc5b* |
| 7 | Cell cycle process | *wee2, ccnb2, ccna1, ccnb1, birc5b, mos, fbxo43* |
| 7 | Regulation of molecular function | *ccnb2, ccna1, ccnb1, gdf9, bmp15, gdf3, pabpn1l* |
| 5 | Response to endogenous stimulus | *cyp11a1, gdf9, bmp15, gdf3, cpeb1b* |
| 5 | Response to chemical | *cyp11a1, gdf9, bmp15, gdf3, cpeb1b* |
| 5 | Regulation of biological quality | *cyp11a1, cth1, rdh10b, tph1a, ca15b* |
| 4 | Cell proliferation | *btg4, ccnb2, ccna1, ccnb1* |
| 4 | Catabolic process | cth1, zgc:171517, nots, map1lc3c |
| 4 | Anatomical structure morphogenesis | *gdf3, cyp11a1, hsd3b2, birc5b* |
| 4 | Sexual reproduction | *bmp15, ccnb1, ca15b, birc5b* |
| 4 | Regulation of signaling | *gdf9, bmp15, gdf3, mos* |
| 4 | Multicellular organism reproduction | *bmp15, ccnb1, ca15b, birc5b* |
| 4 | Multi-organism reproductive process | *bmp15, ccnb1, ca15b, birc5b* |
| 4 | Regulation of response to stimulus | *gdf9, bmp15, gdf3, mos* |
| 4 | Multicellular organismal reproductive process | *bmp15, ccnb1, ca15b, birc5b* |
| 4 | Regulation of developmental process | *tph1a, bmp15, gdf3, bircb5* |
| 4 | Regulation of multicellular organismal process | *tph1a, bmp15, gdf3, bircb5* |
| 4 | Multi-organism process | *bmp15, ccnb1, ca15b, birc5b* |
| 4 | Meiotic cell cycle process | *wee2, mos, fbxo43, birc5b* |
| 4 | Regulation of reproductive process | *wee2, mos, fbxo43, bmp15* |
| 3 | Developmental process involved in reproduction | *bmp15, map1lc3c, cyp11a1* |
| 3 | Cellular component biogenesis | *birc5b, map1lc3c, cyp11a1* |
| 2 | Immune system process | *rag1, tph1a* |
| 2 | Immune system development | *rag1, tph1a* |
| 2 | Response to stress | *map1lc3c, cpeb1b* |
| 2 | Cell adhesion | *cldng, cldnd* |
| 2 | Biological adhesion | *cldng, cldnd* |
| 2 | Hormone metabolic process | *cyp11a1, rdh10b* |

**Table S5.** Ontological analysis of the top-50 downregulated genes in the transcriptomic study. p-value <0.05.

| N | High level GO category | Genes |
|---|---|---|
| 2 | Anatomical structure formation involved in morphogenesis | *gdf3, birc5b* |
| 2 | Localization | *ca15b, birc5b* |

**Table S6.** Confirmation by qPCR of selected DEGs identified in the transcriptome analysis.

| Gene | Gene ID (GeneBank) | Transcriptome | | qPCR | |
|---|---|---|---|---|---|
| | | Fold change | p-value | Fold Change ($2^{-\Delta\Delta Ct}$) | p-value |
| *tuba7l* | 431777 | 49.901 | 7.229E-25 | 12.923 | 0.003 |
| *irg1l* | 562007 | 35.405 | 1.458E-23 | 14.015 | 0.013 |
| *sycp3* | 678602 | 12.110 | 2.602E-21 | 10.938 | 0.009 |
| *stAR* | 63999 | 11.570 | 3.858E-06 | 4.256 | 0.002 |
| *amh* | 493624 | 6.883 | 1.623E-20 | 4.133 | 0.001 |
| *dmrt1* | 402923 | 6.614 | 2.220E-09 | 15.940 | 0.005 |
| *dkk1a* | 799377 | 4.649 | 6.161E-11 | 3.759 | 0.001 |
| *plpp4* | 100334794 | -219.745 | 5.052E-04 | 0.098 | 0.024 |
| *grik3* | 100334689 | -189.490 | 1.818E-04 | 2.763 | 0.066 |
| *cyp11a1* | 80374 | -7.005 | 2.290E-04 | 0.038 | 0.001 |
| *lef1* | 30701 | -5.079 | 1.655E-06 | 0.093 | 0.001 |
| *ctnnbip1* | 58117 | -3.630 | 1.253E-12 | 0.250 | 0.002 |
| *dvl3a* | 80972 | -2.587 | 8.327E-06 | 0.413 | 0.002 |

Supplemental Figures

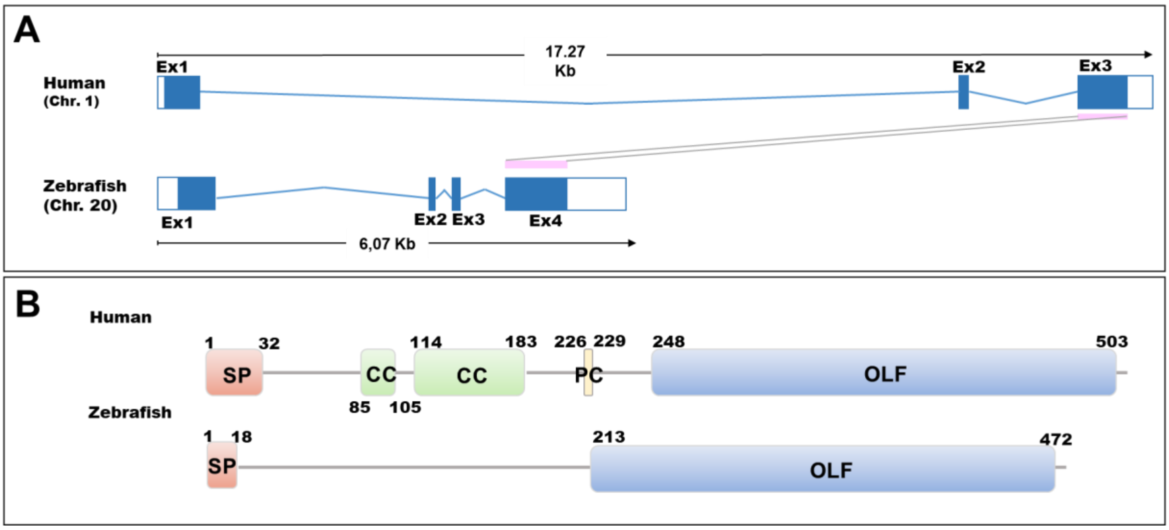


**Figure S1.** Conservation of gene structure and protein domain organization of human and zebrafish myocilin. (A) Genomic alignment of human (ENSG00000034971) and zebrafish (ENSDARG00000021789) genes. The "Ensembl region comparison tool" was used to obtain this image. Pink rectangles represent conserved exons. (B) Human and zebrafish myoc protein domain comparation. Domains are indicated according to the UniProt database (A0A0S2Z421 and Q5F0G5, for the human and zebrafish domains, respectively; https://www.uniprot.org/uniprot). The numbers correspond to amino acid positions. SP: Signal peptide. CC: Coiled coils. OLF: Olfactomedin domain. PC: Proteolytic cleavage site.

```
human            MRFFCARCCSFGPEMPAVQLLLLACLVWD-------VGARTAQLRKANDQSGRCQYTFSV    53
zebrafish        --------------------MWFLAVLWISSLLMGSQVQSSANLRRANAGNGRCQYTFMV    40
                                     : : .::*            :*:**:**  .******* *

human            ASPNESSCPEQSQAMSVIHNLQRDSSTQRLDLEATKARLSSLESLLHQLTLDQAAR--PQ    111
zebrafish        DSPTEASCPSPGSTPEM---------------EALMSRLGLLEALVARLVGGEAMPESSQ    85
                  **.*:***. ..: .:               **  :**. **:*: :*. .:*     *

human            ETQEGLQRELGTLRRERDQLETQTRELETAYSNLLRDKSVLEEEKKRLRQENENLARRLE    171
zebrafish        SSGSGL---------------------QDSYNQVMGENAQLKREKQRLDRQVQDLQQRME    124
                 .: .**                     : :*.::: ::: *:.**:** :: ::* :*:*

human            SSSQEVARLRRGQCPQTRDTA-----RAVPPG--------------------SREVSTWN    206
zebrafish        ELRQEAERLRSRPCMQQTSSRVPQKDNSFRPGSGHVPSNLASRPGNPQEDKSSLRDPAWQ    184
                 .  **. ***   * *  .:      .:. **                    * .  :*:

human            LDTLAFQELKSELTEVPASRILKESPSGYLRSGEGDTGCGELVWVGEPLTLRTAETITGK    266
zebrafish        YSNPGYQELTAVVTEVTAPN---------QDGPADISGCGDLVWVENPEVHRKADSIAGK    235
                  .. .:***.: :*** * .           .  . :***:**** :* . *.*::*:**

human            YGVWMRDPKPTYPYTQETTWRIDTVGTDVRQVFEYDLISQFMQGYPSKVHILPRPLESTG    326
zebrafish        YGVWMQDPEAKEPYGPDMVWRIDSVGSEVRQLFGYENMDQLTRGFPTKVLLLPESVESTG    295
                 *****:**: . **  : .****:**::***:* *: :.*: :*:*:** :**. :****

human            AVVYSGSLYFQGAESRTVIRYELNTETVKAEKEIPGAGYHGQFPYSWGGYTDIDLAVDEA    386
zebrafish        ATMYKGSLYYQRRLSRTLIRYDLHAESIAARRDLPHAGFHGQFPYSWGGYTDIDLAIDEN    355
                 *.:*.****:*   ***:***:*::*:: *.:::* **:*****************:**

human            GLWVIYSTDEAKGAIVLSKLNPENLELEQTWETNIRKQSVANAFIICGTLYTVSSYTSAD    446
zebrafish        GLWAIYSTNKAKGAIVISQLDPHNLEVKGTWETKIRKTSVANAFMICGKLYTVASYTSPN    415
                 ***.****::******:*:*:*.***:: ****:*** ******:***.****:**** :

human            ATVNFAYDTGTGISKTLTIPFKNRYKYSSMIDYNPLEKKLFAWDNLNMVTYDIKLSKM-     504
zebrafish        TTVNYMFDTATSQGKAISVPFKNRYRYNSMVDYNSAKRKLYAWDNYYMVSYSVRLGKQE     474
                 :***: :**.*. .*::::******:*.**:***  ::**:****  **:*.::*.*
```

Figure S2. Amino acid sequence comparison of Human and Zebrafish myocilin. The alignment the human (UniProt accession number A0A0S2Z421) and zebrafish (UniProt accession number Q5F0G5) proteins was carried out with ClustalW (https://embnet.vital-it.ch/software/ClustalW.html). The asterisks indicate the positions where all the amino acids are identical, two vertical dots show amino acids with similar chemical properties and one dot denotes amino acid positions with weak chemical similarity. Orange background: olfactomedin domains.

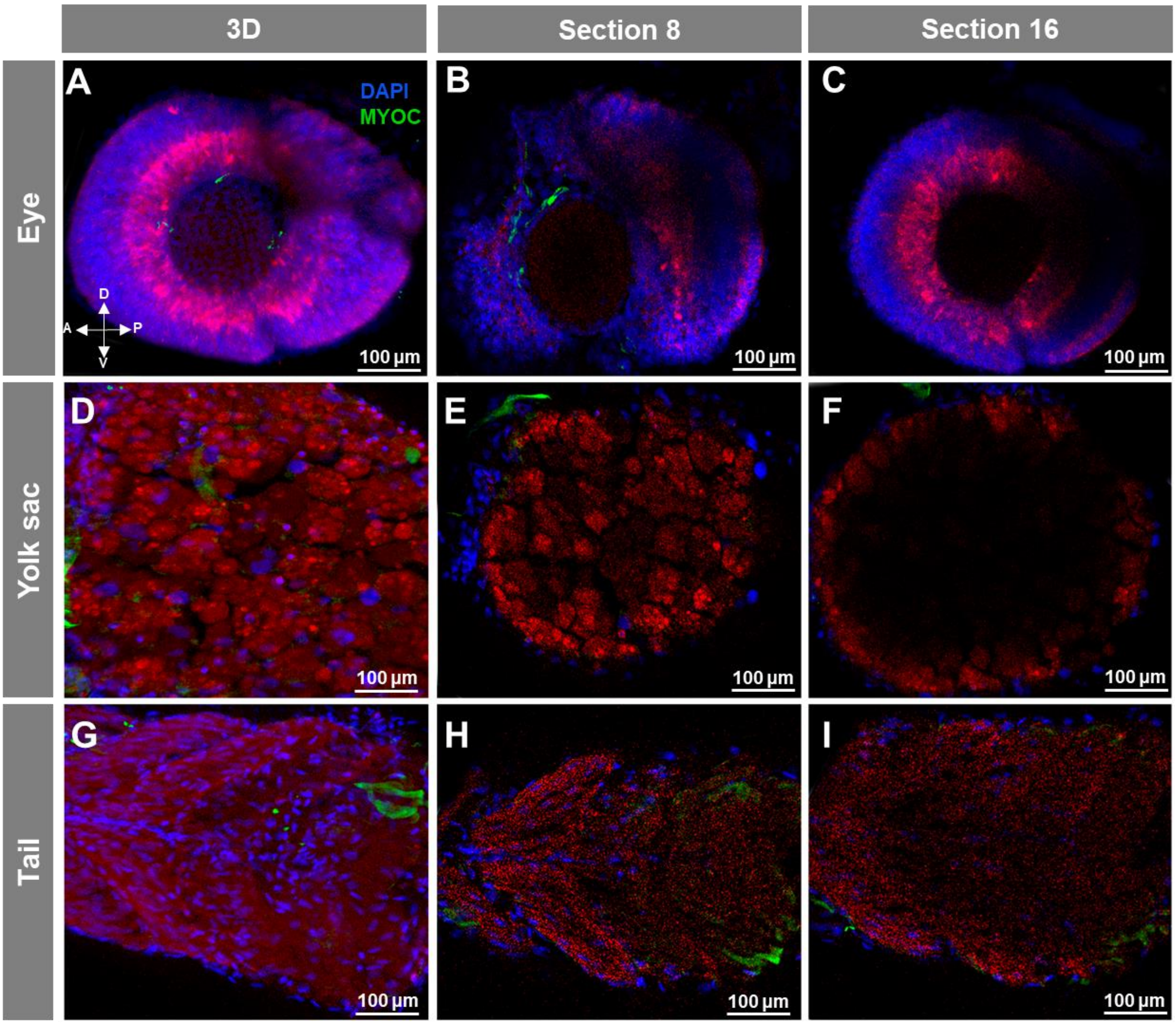


**Figure S3.** Negative controls of fluorescent whole-mount immunohistochemistry of myoc in wild-type zebrafish embryos (96hpf) shown in Figs. 2A-C, 3A-C and 4A-C. Wild-type embryos were incubated with a preimmune TNT antibody and Cy2-conjugated goat anti-chicken IgY secondary antibody. Red: tissue autofluorescence.

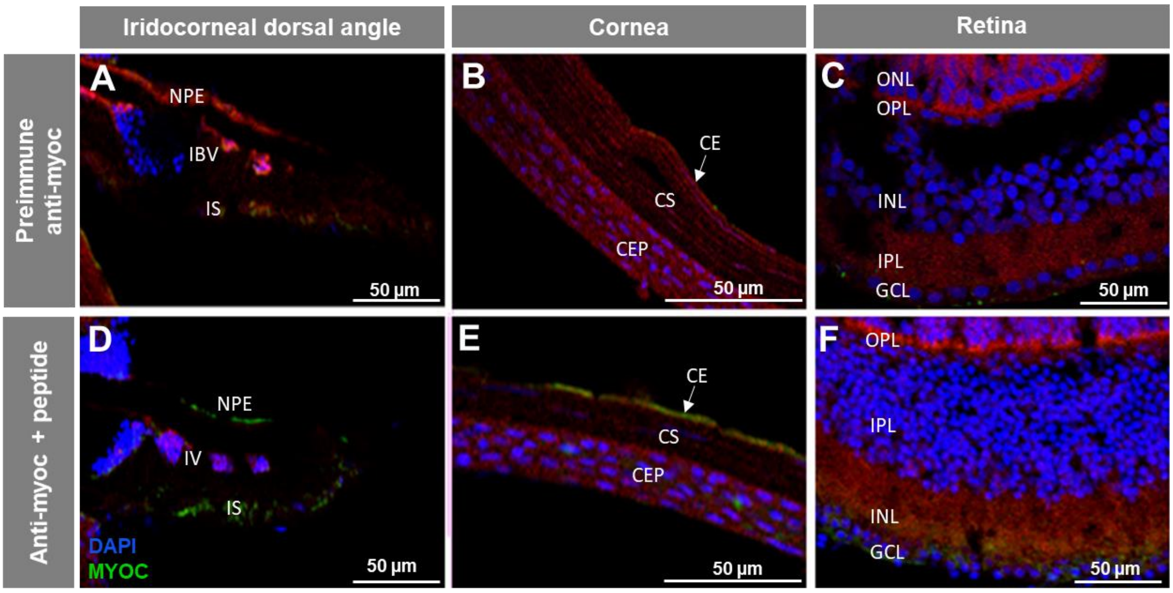


Figure S4. Negative controls of the immunohistochemical myoc analysis in the adult (7 months) wild-type zebrafish eye shown in Figure 5. Controls consisted of tissue sections (14 μm) incubated with a chicken preimmune anti-myocilin antibody (TNT, 1:150), followed by the secondary antibody (A–C). As an additional control, tissue sections were incubated with the antigenic peptide (1:20 antibody:peptide) (D–F). Red signals correspond to tissue autofluorescence. NPE: Non-pigmented ciliary epithelium; IBV: Iris blood vessels; IS: iris stroma. CE: Cornea endothelium; CS: stroma; CE: cornea epithelium GCL: ganglion cell layer; IPL: inner plexiform layer; INL; inner nuclear layer; OPL: outer plexiform layer; ONL; outer nuclear layer; PHL: photoreceptor layer.

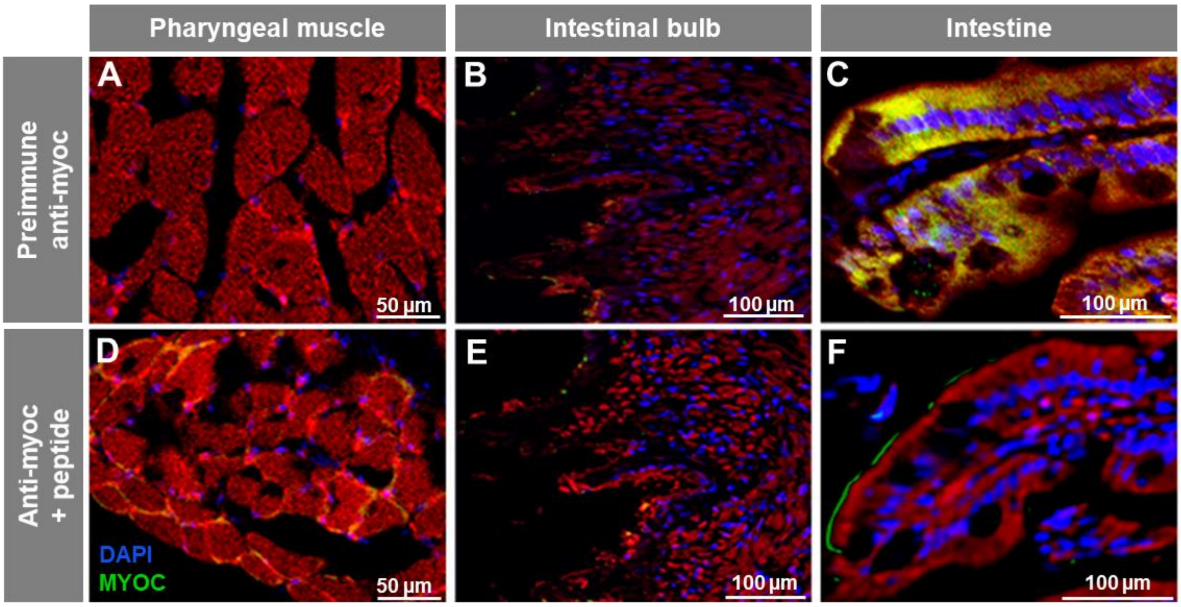


Figure S5. Negative controls of the Immunohistochemistry of myocilin in non-ocular tissues of adult wild-type zebrafish shown in Figures 6A-F. Tissue sections (14 µm) were incubated with the preimmune anti-myocilin antibody (TNT, 1:150), followed by incubation with a Cy2-conjugate goat anti-chicken IgY secondary antibody (1:1000) (A–C). As a competitive control, tissue sections were incubated with a chicken anti-TNT primary antibody (1:150) in the presence of the antigenic peptide (1:10 antibody:peptide). A Cy2-conjugate goat anti-chicken IgY (1:1000) was used as secondary antibody (D–F). Red signal: tissue autofluorescence.

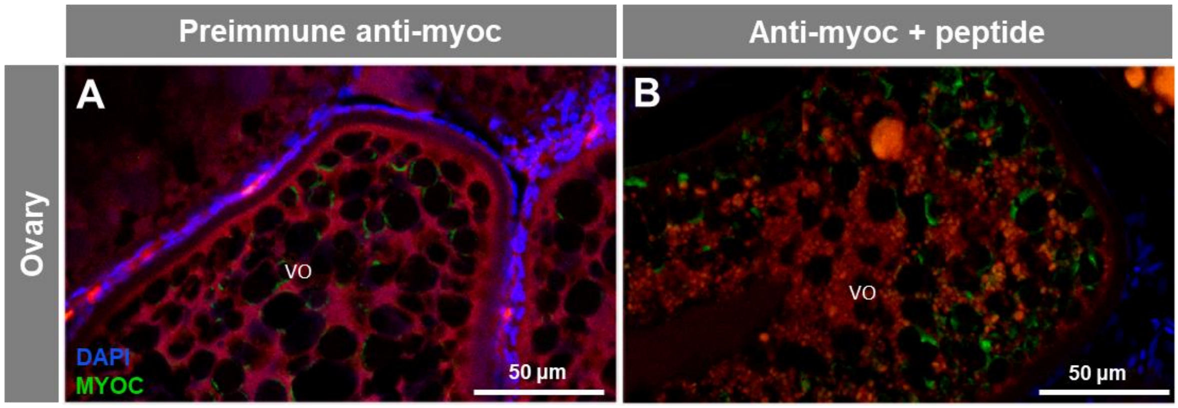


**Figure S6.** Negative control of the Immunohistochemistry of myocilin in the ovary of adult wild-type zebrafish shown in Figures 7A,B. Ovary sections (14 μm) of adult wild-type zebrafish were incubated with the preimmune anti-myocilin antibody (TNT, 1:150) (A). A competitive assay using the antigenic peptide (1:10 antibody:peptide) was carried out as an additional control (B). Red signals correspond to tissue autofluorescence. VO: vitellogenic oocyte.

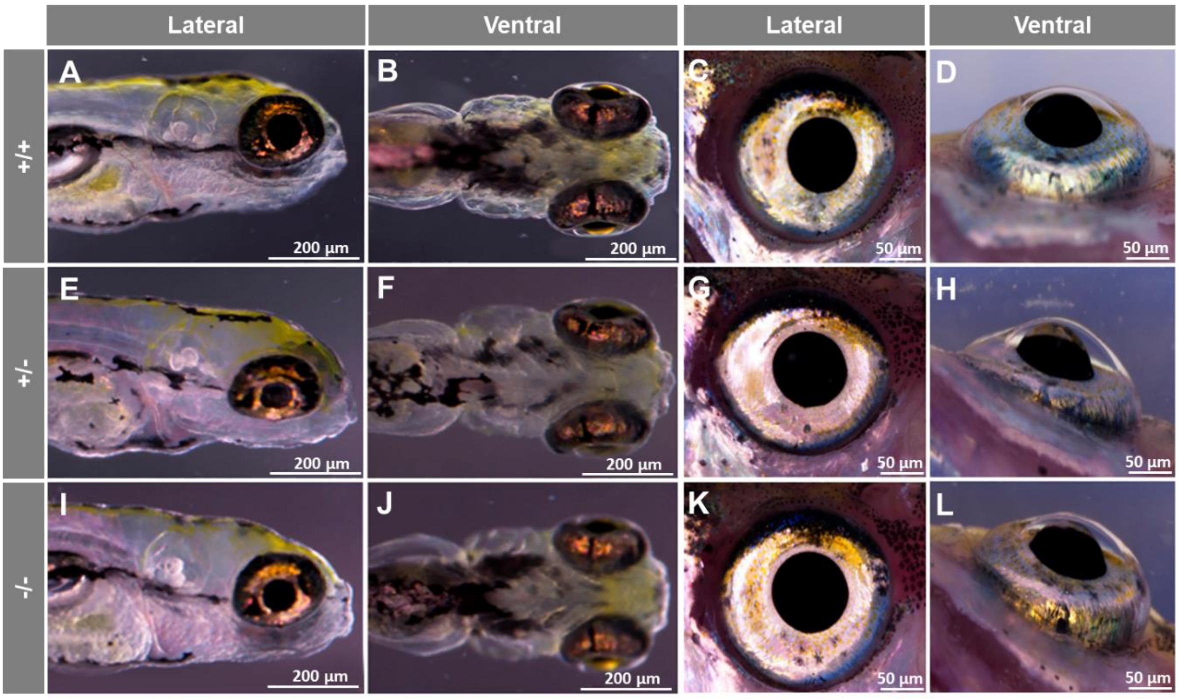


Figure S7. Head and eye macroscopic phenotypes in mutant *myoc* embryo and adult zebrafish. Heterozygous *myoc* mutants were inbred and the progeny was genotyped by PAGE and observed at 96 hpf using a Nikon SMZ18 (A, B, E, F, I, and J). Part of the progeny was raised to adulthood (7 months), genotyped and photographed as described (C, D, G, H, K, and L). No significant phenotypic differences were observed between the different genotypes. The images are representative of ten individuals of each stage and genotype. +/+: wild-type; -/-: heterozygous; -/-: myoc KO.

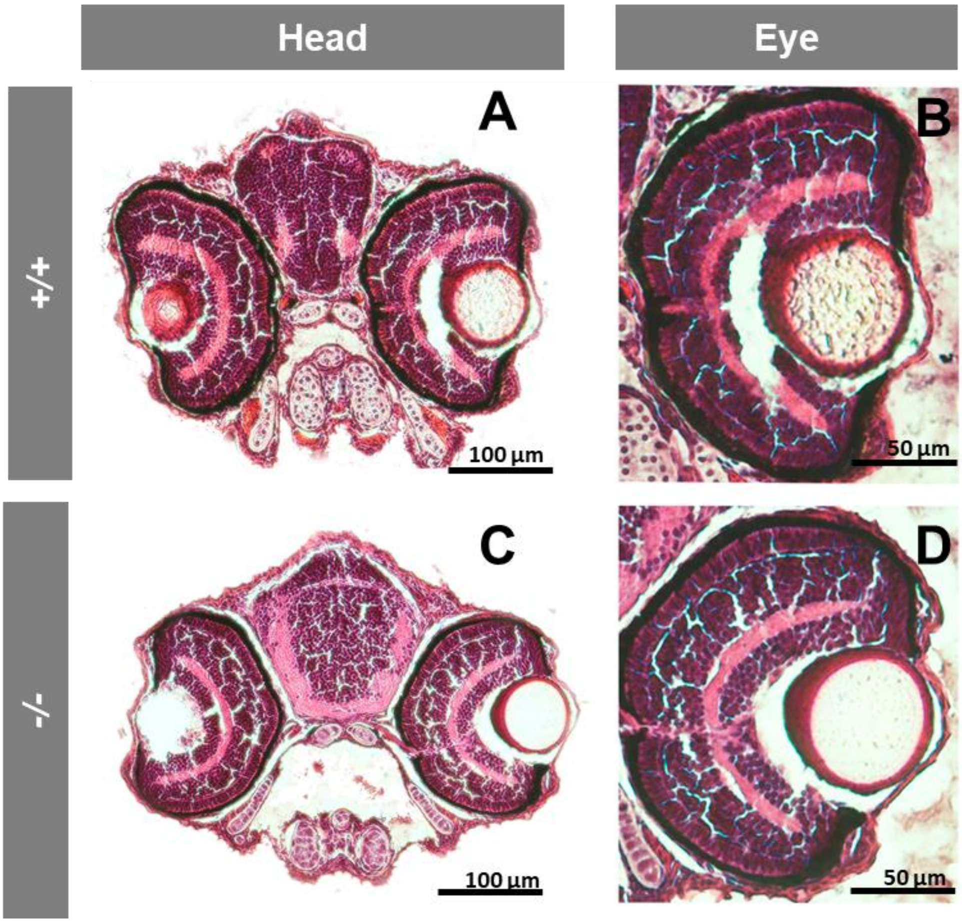


Figure S8. Microscopic head and eye phenotypes in mutant *myoc* zebrafish embryos (96 hpf). The embryos were obtained as described in Figure S7. Tissue sections were stained with hematoxylin-eosin. No significant phenotypic differences were observed between +/+ and -/- embryos. The images are representative of three individuals of each genotype. +/+: wild-type; -/-: myoc KO.

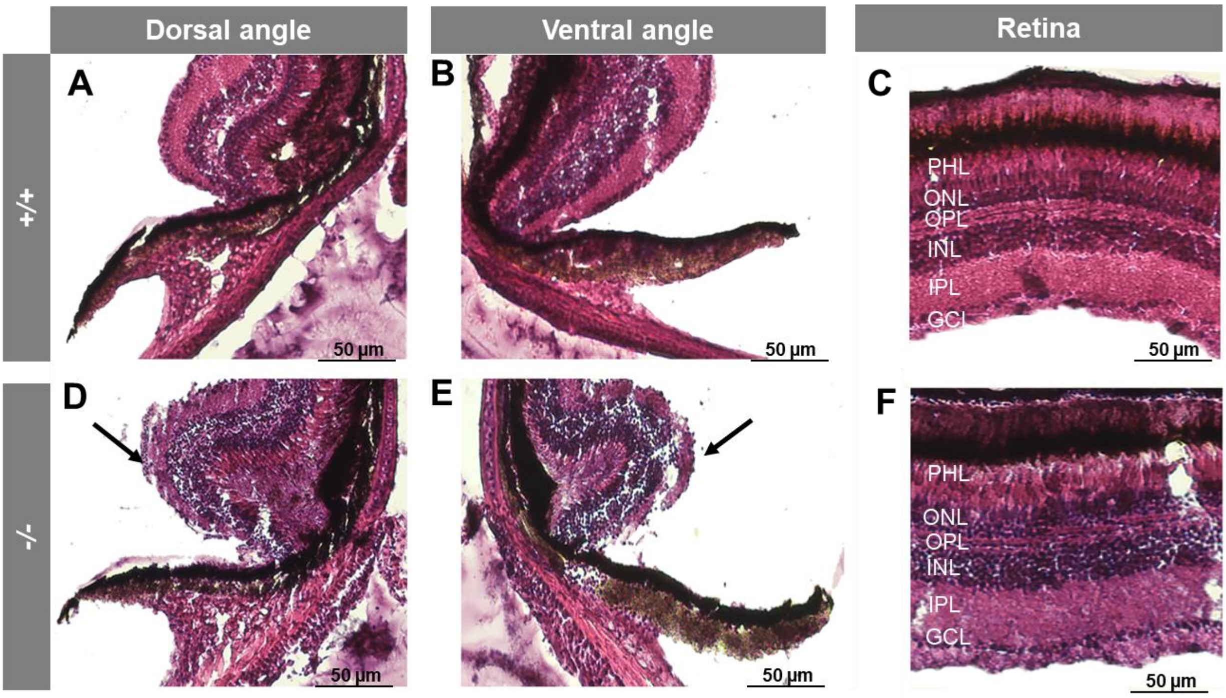


**Figure S9.** Histology of ocular anterior segment and retina from adult mutant *myoc* zebrafish. The fishes were obtained as described in Figure S7. Tissue sections were stained with hematoxylin-eosin. No significant phenotypic differences were observed between +/+ and -/- embryos. An apparently increased folding of the anterior retina was observed in -/- animals (arrow), but it was also detected in some +/+ zebrafish. The images are representative of three individuals of each genotype. +/+: wild-type; -/-: myoc KO. GCL: ganglion cell layer; IPL: inner plexiform layer; INL: inner nuclear layer; IPE: iris pigment epithelium; OPL: outer plexiform layer; ONL; outer nuclear layer; RPE: retinal pigment epithelium.

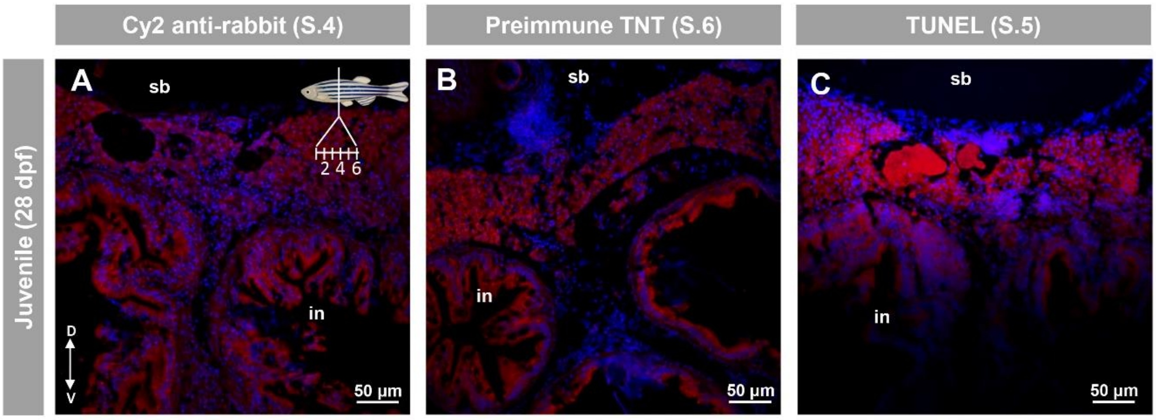


**Figure S10.** Negative controls of immunohistochemistry and TUNEL assay of the immature (28 dpf) zebrafish gonad shown in Figure 9. As a negative control of the anti-vasa antibody used in Figures 9A,B,E and F, tissue section four from wild-type zebrafish was incubated with a Cy2-conjugate goat anti-chicken IgG secondary antibody (1:1000) (A). Tissue section six was treated with an anti-myocilin preimmune antibody (TNT, 1:150) followed by incubation with a Cy2-conjugate goat anti-rabbit IgG secondary antibody (1:1000), as control of the immunohistochemistry shown in Figures 9C,O (B). Tissue section five was treated with only the terminal dUTP nick-end labeling solution a control of the TUNEL assay shown in Fig 9D,H (C). Blue and red signals correspond to DAPI nuclear staining and tissue autofluorescence, respectively. G: gonad. I: intestine. SB: swimbladder. The insert in (A) show the position of the different tissue sections. The vertical double arrow in (A) indicate de dorsovental axis (D: dorsal; V: ventral). The images are representative of the results observed in four larvae.

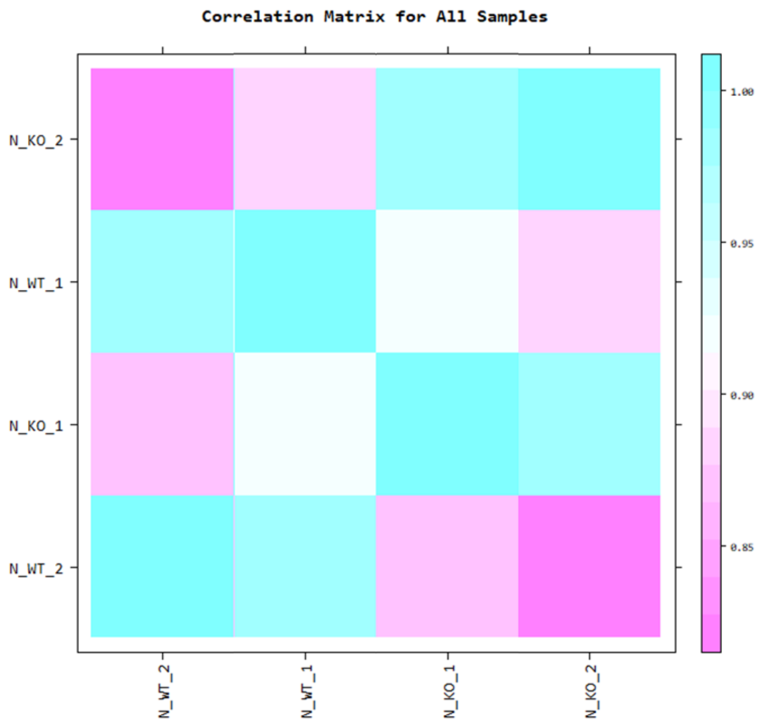


**Figure S11.** Correlation matrix showing the similarity between RNAseq replicas (male *myoc* KO vs. male wild type). The similarity between samples is obtained through Pearson's coefficient of sample's normalized value ($-1 \leq r \leq 1$). The closer the value is to 1, the more similar the samples are.

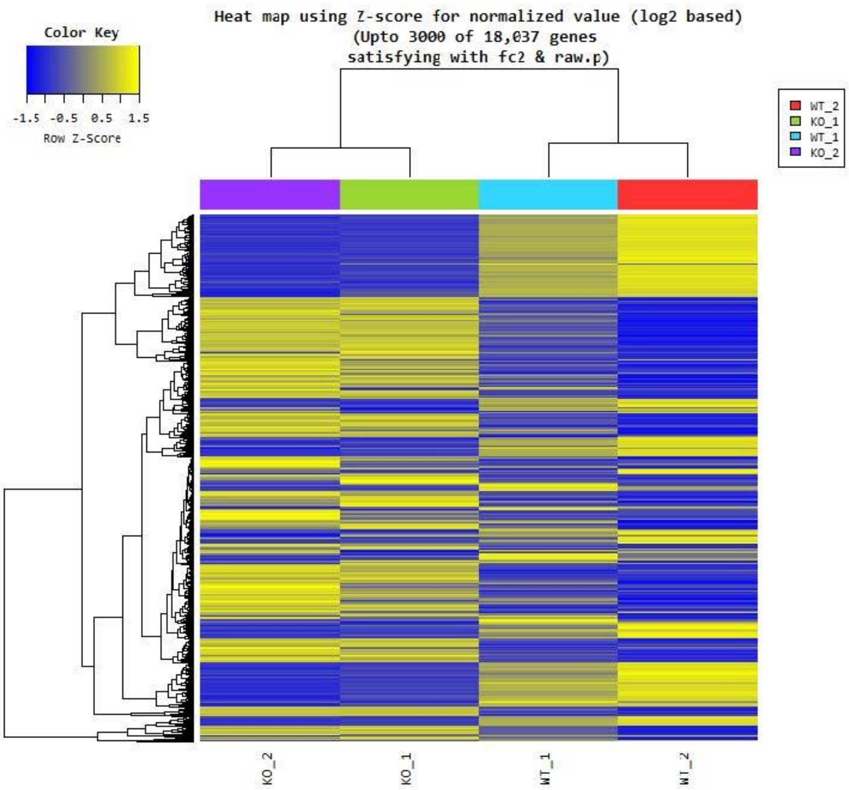


**Figure S12.** Heatmap of hierarchical clustering analysis of significant DEG and replicas (male *myoc* KO vs. male wild type). Clustering analysis was done using Euclidean distance and complete linkage as a measure of similarity. The results show 3000 DEGs which satisfied a fold change > 2 or < -2 and a raw p value < 0.05.

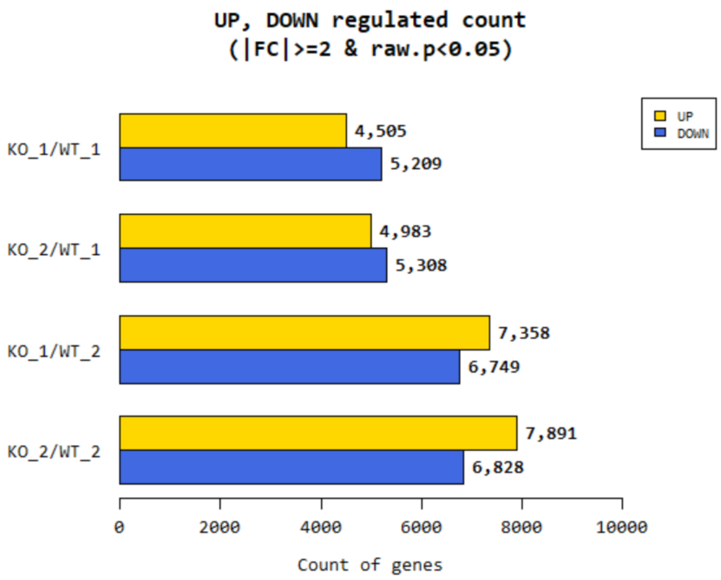


**Figure S13.** Number of significant up and down regulated genes based on fold change of comparison pairs (male *myoc* KO vs. male wild type). The results show DEGs which satisfied a fold change > 2 or < -2 and a raw p value < 0.05.

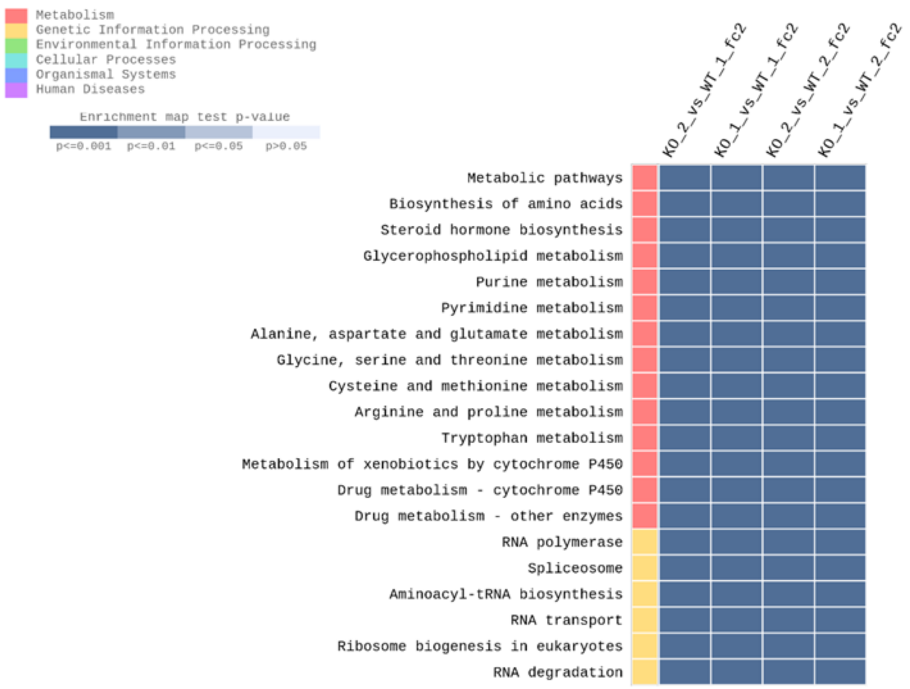


**Figure S14.** Top-20 KEGG pathways differentially expressed in *myoc* knockout male zebrafish in the four comparisons (KO1 vs. WT1, KO1 vs. WT2, KO2 vs. WT1 and KO2 vs. WT2). The pathways were identified by KEEG enrichment analysis. The enrichment map is colored by the gradient level of the raw p value from the modified Fisher's exact test. A raw p-value < 0.05 means that the pathway has been significantly enriched.

Supplementary Video Captions

**Video S1:** Two-dimensional confocal image z-stacks corresponding to whole-mount immunohistochemical detection of myocilin in the eye of 96 hpf-wild-type zebrafish embryos show in Figure 2.

**Video S2:** Two-dimensional confocal image z-stacks corresponding to whole-mount immunohistochemical detection of myocilin in the yolk of 96 hpf-wild-type zebrafish embryos show in Figure 3.
